# *Arabidopsis thaliana egy2* mutants display altered expression level of genes encoding crucial photosystem II proteins

**DOI:** 10.1101/286948

**Authors:** Małgorzata Adamiec, Lucyna Misztal, Ewa Kosicka, Ewelina Paluch-Lubawa, Robert Luciński

## Abstract

EGY2 is a zinc – containing, intramembrane protease, located in the thylakoid membrane. It is consider to be involved in the regulated intramembrane proteolysis - a mechanism leading to activation of membrane-anchored transcription factors through proteolytic cleavage, which causes them to be released from the membrane. The physiological functions of EGY2 in chloroplasts remains poorly understood. To answer the question what is the significance of EGY2 in chloroplast functioning two T-DNA insertion lines devoid of EGY2 protein were obtained and the mutants phenotype and photosystem II parameters were analyzed. Chlorophyll fluorescence measurements revealed that the lack of EGY2 protease caused changes in non-photochemical quenching (NPQ) and minimum fluorescence yield (F_0_) as well as higher sensitivity of photosystem II (PSII) to photoinhibition. Further immunoblot analysis revealed significant changes in the accumulation levels of the three chloroplast-encoded PSII core apoproteins: PsbA (D1) and PsbD (D2) forming the PSII reaction centre and PsbC – a protein component of CP43, a part of inner PSII antennae. The accumulation level of nuclear-encoded proteins Lhcb1-3 - a components of the major light-harvesting complex II (LHCII) as well as proteins forming minor peripheral antennae complexes, namely Lhcb4 (CP29), Lhcb5 (CP26), and Lhcb6 (CP24) remain, however, unchanged. The lack of EGY2 led to a significant increase in the level of PsbA (D1) with simultaneous decrease in accumulation levels of PsbC (CP43) and PsbD (D2). To test the hypothesis that the observed changes in the abundance of chloroplast-encoded proteins are a consequence of changes in gene expression levels, real-time PCR was performed. The obtained results shown that *egy2* mutants display an increased expression of *PSBA* and reduction in the *PSBD* and *PSBC* genes. Simultaneously pTAC10, pTAC16 and FLN1 proteins were found to accumulate in thylakoid membranes of analyzed mutant lines. These proteins interact with core complex of plastid encoded RNA polymerase and may be involved in the regulation of chloroplast gene expression.

## Introduction

Regulated intramembrane proteolysis (RIP) is a mechanism that regulates gene expression at the transcriptional level. This process involves the activation of membrane-anchored transcription regulators through proteolytic cleavage, which causes them to be released from the membrane. The proteases participating in this process perform proteolytic cleavage within the cell membrane and are referred to as intramembrane cleaving proteases (I-CLiPs) (Weihofen et al., 2002, Koonin et al., 2003, Kinch et al., 2006). Site-2 proteases (S2P), zinc-containing metalloproteases that are able to perform proteolytic cleavage within a membrane, belong to the CLiPs family (Kinch et al., 2006). Representatives of this unusual and relatively recently discovered family of proteases are present in all living organisms. However, knowledge concerning their substrates, mechanisms of action and physiological roles remains rather limited. To date, several substrates of human and bacterial representatives of the S2P family have been identified. The first S2P substrate to be identified was human sterol regulatory element binding protein (SREBP), a transcription factor that is involved in the regulation of sterol and fatty acid synthesis (Brown and Goldstein, 1997). Other substrates also, although involved in very diverse physiological processes like responses to the accumulation of unfolded proteins in the endoplasmic reticulum (Ye et al., 2000; Kondo et al., 2005; Zhang et al., 2006), extracytoplasmic stress responses (Schobel et al., 2004) or the regulation of cell division (Bramkamp et al., 2006) have been identified as transcription factors.

A very little is known about the physiological functions of S2P proteases in plants. Five genes encoding S2P homologues with potential proteolytic activity were identified in the *Arabidopsis thaliana* genome, and the activity of three of them was confirmed experimentally (Chen et al., 2005; Che et al., 2010; Chen et al., 2012). The localization of all of the proteases was confirmed experimentally. Four of them were found in chloroplasts and one in the Golgi membrane (for a review see Adamiec et al., 2017). The only known substrate of plant S2P proteases is bZIP17, which is cleaved by the S2P protease encoded by *At4g20310*. bZIP17 was found to control the expression levels of the transcription factor AtHB7 and the protein phosphatases HAB1, HAB2, HAI1 and AHG3, which are negative regulators of abscisic acid signalling (Zhou et al., 2015) and participate in the response to salt stress (Liu et al., 2007). It was also suggested that the S2P protease encoded by *At4g20310* may be involved in processing the transcription factor bZIP28 (Gao et al., 2008). There is, however, no direct experimental evidence confirming this prediction. The ARASP protease has been demonstrated to be essential for plant development and chloroplast biogenesis; however, deeper insight into ARASP-mediated events is lacking (Bölter et al., 2006). ARASP is closely co-expressed with another S2P homologue, S2P2 (Aoki et al., 2016). There is, however, no other evidence indicating cooperation between these proteases, and the physiological role of S2P2 remains unknown. The EGY1 protease is another *A. thaliana* S2P that is involved in chloroplast development (Chen et al., 2012). Additionally, in *egy1 A. thaliana* mutants, a deficiency in ethylene-induced gravitropism was detected (Guo et al., 2008). However, in this case, the mechanisms leading to phenotypic effects were not described. It seems possible that at least some of chloroplast located S2P may be involved in regulation of chloroplast gene expression. In higher plants transcription of chloroplast genes is carried out by two distinct types of RNA polymerases: PEP (plastid – encoded RNA polymerase) and NEP (nuclear-encoded plastid RNA polymerase). NEP is monomeric, phage-type RNA polymerase represented in *Arabidopsis thaliana* and other eudicots chloroplasts by RPOTp and RPOTmp (Elis and Hartley, 1971; Chang et al., 1999; Lagen et al., 2002). PEP, in turn, is holoenzyme functionally similar to *E. coli* RNA polymerase. The core of the enzyme is composed of the sigma subunit and four additional subunits: α, β, β’and β” (Hajdukiewicz et al., 1997). The sigma subunit is responsible for promoter recognition and initiation of transcription. In *Arabidopsis thaliana* six genes encoding sigma factors SIG1-SIG6 are present (for a review see Lysenko, 2007, Lerbs-Mache, 2011). The PEP core complex interacts with many additional proteins, called PEP-associated proteins (PAPs), like DNA gyrase, DNA polymerase, three ribosomal proteins (L12-A, S3 and L29), phosphofructokinase–B type enzymes PFKB1 and PFKB2 or Fe-dependent superoxide dismutases (Fe-SODs). Recently additional PAPs, named plastid transcriptionaly active chromosome proteins (pTAC) were identified. These proteins were proven to be located within chloroplast membranes (Hess and Borner, 1999). In *Arabidopsis thaliana* genome 18 membrane attached pTAC proteins were identified, however their function and mechanism of action are poorly investigated (Pfalz et al., 2006). The majority of chloroplast genes are transcribed mainly by PEP or both PEP and NEP (Allison et al., 1996; Hajdukiewicz et al., 1997)

The EGY2 protease is also present in chloroplast membranes, and its proteolytic activity was demonstrated experimentally (Chen et al., 2012). The EGY2 protease contains 556 amino acid (aa) residues, and the number of predicted transmembrane domains varies from 5 to 8 (Schwacke et al., 2003). The zinc atom is probably coordinated by and located in the hydrophobic region formed by H^324^, H^328^ and D^460^, and the conserved structural region of the protein extends from A^375^ to G^380^ (Lamesch et al., 2011). The substrates of EGY2 remain unknown, and the physiological function of the protease has been poorly investigated. Although *egy2* knockout mutants do not display a clearly visible phenotype, EGY2 was found to play a role in hypocotyl elongation in *A. thaliana* (Chen et al., 2012). The EGY2 protease was also demonstrated to be involved in fatty acid biosynthesis. In *egy2* mutant seedlings, a decrease in overall fatty acid content was observed as well as reduced accumulation of acyl carrier protein 1 and of CAC2 and BCCP1, two subunits of plastidic acetyl-coenzyme A carboxylase (ACCase) (Chen et al., 2012).

To gain deeper insight into the role of the EGY2 protease in chloroplasts, we analysed changes in the accumulation of apoproteins that are encoded in the nuclear and chloroplast genomes and that are crucial for PSII structure. In this study, we showed that lack of EGY2 protease influences the levels of the chloroplast-encoded apoproteins PsbA (D1), PsbD (D2) and PsbC (CP43) within the PSII core centre and that EGY2-mediated regulation of these proteins occurs at the transcriptional level.

## Material and methods

### Plant material and growth conditions

Wild-type (WT) and mutant *Arabidopsis thaliana* (L.) Heynh (ecotype Columbia) plants were grown on sphagnum peat moss and wood pulp in 42-mm Jiffy peat pellets (AgroWit, Przylep, Poland) under long-day conditions (16 h of light and 8 h of darkness) at an irradiance of 110 µmol m^-2^ s^-1^, a constant temperature of 22°C and a relative humidity of 70 %.

*A. thaliana* seeds with a T-DNA insertion in the *EGY2* gene (*At5g05740*) were obtained from NASC (Nottingham Arabidopsis Stock Centre, Nottingham, UK). Two mutant lines were analysed: SALK_028514C, which was previously described by Chen et al. (2012) as *egy2-3*, and SALK_093297C, which was not described earlier. We decided to maintain the nomenclature used by Chen et al. (2012) and therefore, we refer SALK_028514C as *egy2-3*, and SALK_093297C as *egy2-5*. Homozygosity of the T-DNA insertion within the analysed gene was confirmed by PCR.

For both lines, the following primers were used:

forward: 5’-GGAACCAGAAGGCAATGATGATG-3’

reverse: 5’-AACCAGCAGCAAACCATTCAG-3’

T-DNA insertion (LB): 5’-CCCTATCTCGGGCTATTCTTTTG-’3.

All measurements and analysis were performed in three biological replicates, on plants in developmental phase 6.0 according to the BBCH scale (Boyes et al., 2001). Thirty plants from each variant (WT, *egy2-3* and *egy2-5*) were measured in each replicate.

### Nucleic acid analysis

DNA was isolated using the Phire Plant Direct PCR Master Mix (Thermo Fisher Scientific, Waltham, MA, USA). Total RNA for real-time PCR analysis was isolated using the GeneMATRIX Universal RNA Purification Kit (EUR_x_^®^, Poland). Reverse transcription was performed using the RevertAid H Minus First Strand cDNA Synthesis Kit (K1631, Thermo Fisher Scientific) with random hexamers as primers and 5 µg of total RNA from the WT and *egy2* mutant lines. Real-time PCR was performed using the Rotor-Gene 6000 real-time rotary analyser (Corbett, Life Science Technology, Australia) and Maxima™ SYBR Green/ROX qPCR Master Mix (Thermo Fisher). One microliter of cDNA was used for the qPCR reaction, and two cycling conditions were applied:

A. for the *PSBA, PSBD* and *PSBC* genes: 94°C for 20 s, 55°C for 30 s and 70°C for 30 s (40 cycles)
B. for the *PSBI* gene: 94°C for 20 s, 55°C for 30 s and 65°C for 30 s (40 cycles).

All primers were tested for nonspecific amplification and primer-dimer formation by melting curve analysis. As the reference, a gene encoding chloroplast ribosomal protein L2 was used. For each sample, six biological repetitions were performed, each in two technical repetitions. The primers used were as follows:

*PSBA*

F: 5’-ATACAACGGCGGTCCTTATGAAC-3’

R: 5’-CAAGGACGCATACCCAGACGG-3’

*PSBI*

F: 5’-ATGCTTACTCTCAAACTTT-3’

R: 5’-TTATTCTTCACGTCCCGGAT-3’

*PSBD*

F: 5’-GGATGACTGGTTACGGAGGG-3’

R: 5’-GAACCAACCCCCTAAAGCGA-3’

*PSBC*

F: 5’-GCTCCTTTAGGTTCGTTAAATTCTG-3’

R: 5’-AGAACAAAATGAGAGGTAGATAACC-3’

Gene encoding ribosomal protein L2 (AtCg00830)

F: 5’-ATGGAGGTGGTGAAGGGAGGG-3’

R: 5’-TTTTTCCTTTTTCTAGTTCTTCTTCC-3’

The amplification efficiency and the expression levels were calculated using Miner (http://ewindup.info/miner/) (Zhao and Fernald, 2005).

### Protein analysis

#### Antibodies

Anti-EGY2 specific polyclonal antibodies were produced in rabbits by Agrisera AB (Vannas, Sweden) using the highly purified N-terminal region (aa 51-225) of EGY2 from *Arabidopsis.*

Anti-Lhcb1-6, anti–PsbA, anti-PsbC and anti -PsbD antibodies were purchased from Agrisera AB (Vannas, Sweden).

#### Isolation of proteins and determination of protein concentration

Total protein was isolated from 100 mg of *A. thaliana* leaf tissue using Protein Extraction Buffer (PEB, Vannas, Agrisera). The concentration of the extracted protein was determined using the modified Lowry method with a Lowry DC kit (Bio-Rad, Hercules, CA, USA).

#### SDS-PAGE and immunoblotting

SDS-PAGE was performed according to Laemmli (Laemmli, 1970) using 12 % (w/v) polyacrylamide gels. After electrophoresis, the proteins were transferred to PVDF membranes (Bio-Rad USA) and incubated with primary antibodies against PsbA, PsbC, PsbD and Lhcb1-6 after blocking of the membranes with 4 % (w/v) BSA (BioShop, Burlington, Canada). After incubation with secondary antibodies (Agrisera, Vannas, Sweden), the relevant bands were visualized on X-ray film using an RTG Optimax X-ray Film Processor (Protec GmbH, Oberstenfeld, Germany) following a 5-minute incubation of the PVDF membrane with Clarity Western ECL Substrate (Bio-Rad, Hercules, CA, USA). Quantification of the immunostained bands was performed using GelixOne software (Biostep GmbH, Jahnsdorf, Germany). For each primary antibody used, the range of linear immunoresponse was checked (Fig. S1). Only blots that showed a linear relationship between the strength of the signal and the amount of protein used were analysed.

#### Isolation of chloroplasts and thylakoid membranes

Chloroplasts were isolated from 20 g of *A. thaliana* leaf tissue using the Sigma Chloroplast Isolation Kit (Sigma-Aldrich, St. Luis, MO, USA). Leaves were homogenized in an ice-cold homogenization buffer with addition of 1 % (v/v) Protease Inhibitor Cocktail (PIC; Sigma-Aldrich, St. Luis, MO, USA) to avoid proteolysis. The homogenate was filtered through Mesh 100 filter and the filtrate was centrifuged at 200 g for 1 min to remove the unbroken cells. The supernatant was then centrifuged for 10 min. at 1500 g to sediment the chloroplasts, which were then resuspended in the homogenization buffer with 1 % (v/v) PIC. To obtain the intact chloroplasts the chloroplast suspension was centrifuged through 40 % (w/v) Percoll for 6 min at 1700 g. To prepare the thylakoid membranes, the pellet of intact chloroplasts was resuspended in the lysis buffer with addition of 1 % (v/v) PIC and centrifuged for 10 min at 12250 g. The thylakoid membranes were collected as a green pellet and used for protein extraction.

#### Protein extraction from the thylakoids, 2D-electrophoresis and protein identification

The thylakoids were homogenized on ice with the EB buffer (Tris-HCl pH 7.5, 25 % (w/v) Sucrose, 5 % glycerol, 10 mM EDTA, 10 mM EGTA, 5 mM KCl, 1 mM DTT) with addition of 0.5 % (w/v) PVPP and 1 % (v/v) PIC to avoid proteolysis. The thylakoid suspension was then centrifuged for 3 min at 600 g and the supernatant was diluted 2-times with water to reach a 12% concentration of sucrose in the EB buffer and centrifuged for 60 min at 100 000 g. The pellet was collected and resuspended in the Tris-HCl buffer (pH 7.5) containing 5 mM EDTA and EGTA and 1 % (v/v) PIC. The protein concentration was measured using Bradford method (Bradford, 1976). Proteins were solubilized in the presence of 2 % (w/v) Brij^®^ 58 (Sigma - Aldrich, St. Luis, MO, USA) for 1 h at 4°C and then precipitated with acetone with 10 % (w/v) TCA and 0.07 % (v/v) β-mercaptoethanol overnight at −20°C. The precipitated proteins pelleted by centrifugation for 15 min at 20 000 g and then washed three times with pure acetone. The obtained protein pellet was resuspended in buffer containing 7 M urea, 3 M thiourea, 2 % (w/v) amidosulfobetaine-14 (ASB-14; Sigma - Aldrich, St. Luis, MO, USA) and 65 mM DTT for 2 h in room temperature with constant gentle shaking and applied for isoelectrofocusing.

Isoelectrofocusing was carried out using the gel strips forming an immobilized pH gradient from 3 to 10 (Bio-Rad, Hercules, CA, USA). Strips were rehydrated overnight at room temperature and the isolectrofocusing was performed at 18°C in the Protean i12 IEF Cell (Bio-Rad, Singapore) for 90 min at 300 V, 90 min at 3500 V, 20000 VHr at 5500 V. After IEF strips were equilibrated according to Kubala et al. (2015) and the separation of proteins according to their molecular mass was performed using denatured electrophoresis in the 12 % (w/v) acrylamide gels with addition of 6 M urea. After electrophoresis the gels were stained with Coomasie Brilliant Blue (CBB) G-250 and photographed with the use of ChemiDoc^™^MP Imaging System (Bio-Rad, Hercules, CA, USA).

Spot detection and image analysis were performed with the PDQuest^™^Advanced 2-D Gel Analysis Software (Bio-Rad, Hercules, CA, USA). Four images representing two independent biological replicates for each *A. thaliana* lines (WT, *egy2-3* and *egy2-5*) were used. The differentially accumulated proteins (P < 0.05) between the WT and both mutant plant lines with a ratio at least 2.0 in absolute value of protein abundance were excised manually under sterile condition and analyzed by liquid chromatography coupled to the mass spectrometer in the Laboratory of Mass Spectrometry, Institute of Biochemistry and Biophysics, Polish Academy of Science (Warsaw, Poland) as previously described Kubala et al. (2015). The raw data were processed using the Mascot Distiller software (ver. 2.4.2.0, MatrixScience) and then the obtained protein masses and fragmentation spectra were matched TAIR (The Arabidopsis Information Resource) filter using the Mascot Daemon engine search. The search parameters were set as follows: trypsin enzyme specificity, peptide mass tolerance ± 20 ppm, fragment mass tolerance ± 0.6 Da, unrestricted protein mass, one missed semiTrypsin cleavage site allowed, fixed cystine alkylation by carbamidomethylation and methionine oxydation set as variable modification.

Only the peptides with the Mascot score exceeding the threshold value corresponding to < 0.05 false positive rate were considered as positively identified.

#### Determination of chlorophyll and carotenoid concentration*s*

The chlorophyll and carotenoid concentrations were measured using a DMSO assay (Hiscox and Israelstam, 1979). The following equations were used to determine the concentrations (µg/ml) of chlorophyll a (Chl *a*), chlorophyll b (Chl *b*) and total carotenoids (C *x+c*), defined as sum of xanthophylls (x) and carotenes (c) (Sumanta et al., 2014).

Chl *a =* 12.47 A_665_ – 3.62 A_649_

Chl *b* = 25.06 A_649_ – 6.5 A_665_

C *x+c* = (1000 A_470_ – 1.29 Chl *a* – 53.78 Chl *b*)/220

#### Chlorophyll fluorescence measurements

Chlorophyll fluorescence measurements were conducted using FMS1 (Photon System Instruments, Brno, Czech Republic) run by Modfluor software. Before each measurement, the leaves were dark-adapted for 30 min. The measurements were performed according to the protocol described by Genty et al. (1989). The minimum fluorescence yield (F_0_) was established at the beginning of measurement. The maximum quantum efficiency of PSII photochemistry (F_v_/F_m_) and quantum efficiency of open centres in the light (F_v_’/F_m_’) was calculated according to Genty et al. (1989). The applied actinic light intensity was equal to the irradiance prior dark-adaptation: 110 µmol m^-2^ s^-1^. The photochemical fluorescence quenching coefficients qP, ΦPSII as well as non-photochemical quenching parameter (NPQ) were calculated according to Maxwell and Johnson (2000). Thirty plants from each variant (WT, *egy2-3* and *egy2-5*) were measured in each replicate.

#### Rosette area measurements

The rosette area was determined from plant photographs using ImageJ 1.50i software (National Institutes of Health, Bethesda, MD) (https://imagej.nih.gov/ij/).

#### Statistical analysis

Differences in the measured parameters were analysed for statistical significance using one-way ANOVA. Means were regarded as significantly different at P < 0.05.

## Results

### EGY2 T-DNA insertion mutants

The function of the EGY2 protease was studied in two commercially available lines with T-DNA insertion mutations. The lines were obtained from the Nottingham Arabidopsis Stock Centre. Lines SALK_028514C (*egy2-3)* and SALK_093297C (*egy2-5)* were chosen for analysis. The line *egy2-3* was previously described by Chen et al. (2012) as the line with single T-DNA insertion located in the fifth exon, however, in our model, the T-DNA insertion is located in the fourth exon (Fig. 1A). The observed difference is related to the splicing form of the investigated gene. The RT-PCR product of gene described by Chen et al. (2012) was 1584 bp (corresponding to 527 aa) in length; according to TAIR (Lamesch et al., 2011). This corresponds to the second splicing form of the *EGY2* gene model. In our RT–PCR analysis, we obtained a 1671-bp long coding sequence (encoding 556 aa); the length of our product is consistent with that of the first splice variant. The *egy2-5* line was not previously described. Our analysis indicated the presence of two T-DNA insertions located in the fifth and sixth exons (Fig. 1A). To verify the number of T-DNA insertions in the *egy2-*5 mutant, PCR was performed using different combinations of primers for both WT and *egy2-5*. As shown in Fig. 1B, no product from the WT DNA was obtained if one of the primers used was a T-DNA (LB) primer. In contrast, the PCR product sizes obtained when the A-LB and B-LB primer pairs were 1200-bp and 400-bp, respectively, indicated the presence of two T-DNA insertions. The absence of the EGY2 protease from both the *egy2-3* and *egy2-5 lines* was confirmed by immunoblot analysis (Fig. 1C).

**Figure 1.**
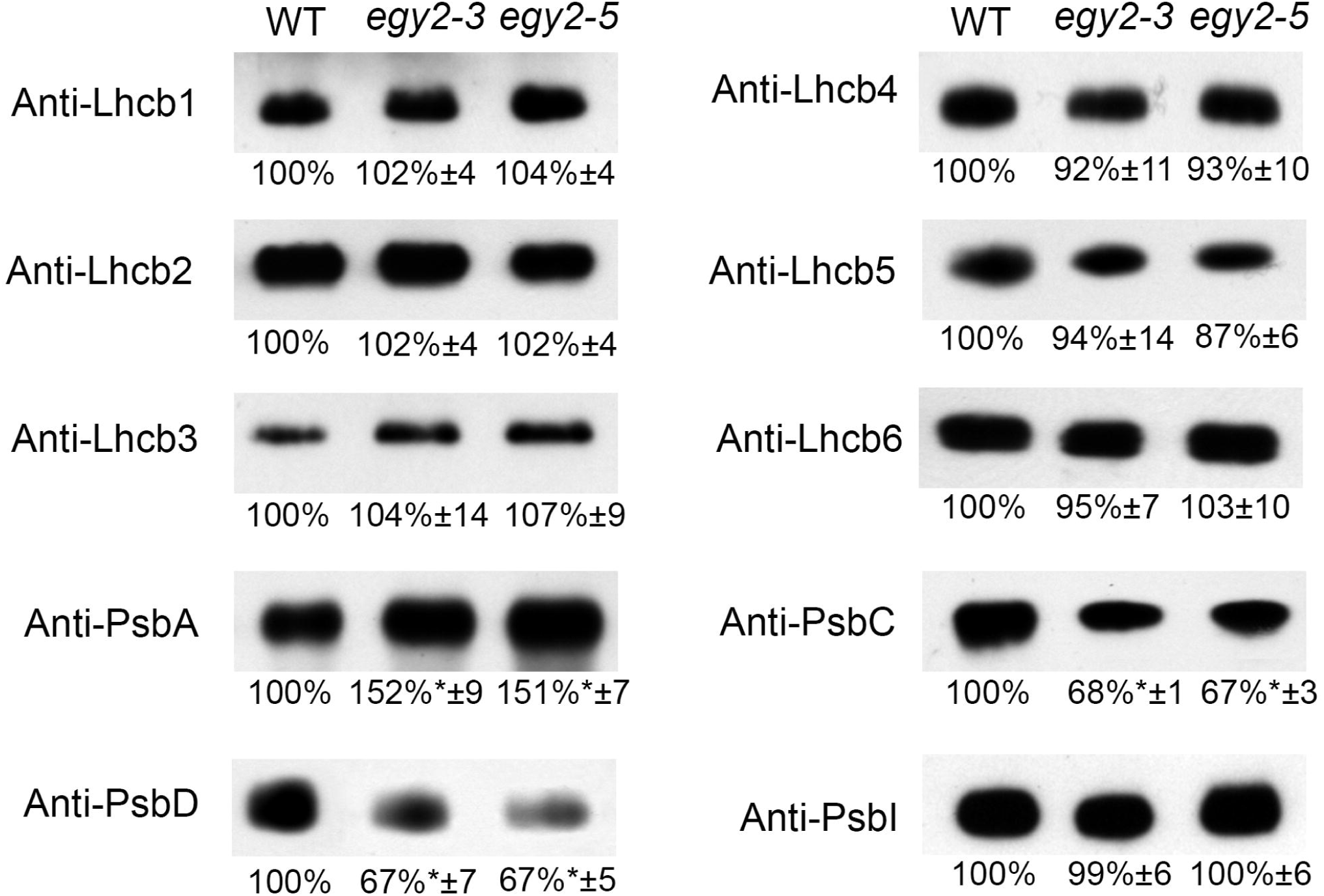
Identification of *egy2* mutants. (A) Schematic diagram of the *Arabidopsis thaliana EGY2* gene. The black boxes represent exons; introns are shown as black lines. The triangles show the locations of T-DNA insertions. The arrows mark the annealing sites of the primers used for PCR analysis. (B) Confirmation of the homozygosity of the *egy2-3* and *egy2-5* mutants. Amplification was performed using the A, B and LB primers as indicated in Fig. 1A. (C) Immunoblot analysis of the EGY2 protease protein in the wild-type (WT) and *egy2-3* and *egy2-5* mutants. Samples of total protein (2, 5 and 10 µg) were resolved by SDS-PAGE, electrotransferred to PVDF membranes and immunostained with the anti-EGY2 antibody.

### Phenotype of *egy2* mutants

The phenotypic analysis revealed no differences between *egy2* mutants and WT plants in leaf or inflorescence emergence or in flower or seed production (data not shown). Both mutant lines, however, displayed significantly larger final rosette areas than those observed in WT plants (Fig. 2). A slight but statistically significant decrease in chlorophyll content was also observed (Tab. 1). This reduction was observed for both chl *a* and chl *b* and was accompanied by a statistically significant reduction in the chl *a*/*b* ratio. A statistically significant reduction in total carotenoid accumulation was also observed (Tab. 1). Chlorophyll fluorescence measurements revealed that the lack of EGY2 protease did not cause statistically significant differences in the F_v_/F_m_ ratio, which describes the maximum quantum yield of PSII photochemistry, or in F_v_’/F_m_’, which indicates the maximum efficiency of PSII photochemistry in the light. The ΦPSII parameter, which determines the quantum efficiency of PSII electron transport in the light, and qP, which is the coefficient of photochemical quenching and is associated with the proportion of open PSII (Maxwell and Johnson, 2000; Murchie and Lawson, 2013), also remained unchanged (Tab. 1). The only observed changes were found in non-photochemical quenching (NPQ), which is thought to be linearly related to heat dissipation (Maxwell and Johnson, 2000), and minimum fluorescence yield (F_0_). Both F_0_ and NPQ were significantly increased in the two mutant lines under normal light conditions (110 µmol m^-2^ s^-1^) compared to their values in WT plants (Tab. 1). To determine whether the lack of EGY2 protease in *egy2* mutants affect the plant’s sensitivity to photoinhibition, changes in the maximum quantum yield of PSII (F_v_/F_m_) were investigated under high light conditions according to our previous work (Luciński et al., 2011). The maximum quantum efficiency of PSII photochemistry (F_v_/F_m_) was measured in plants exposed for 2 h to photoinhibitory irradiance (800 µmol m^-2^ s^-1^); the plants were then illuminated for another 2 h with comfort irradiance (110 µmol m^-2^ s^-1^) to examine the ability of PSII to recover. The initial value of F_v_/F_m_ was similar in the mutant and WT plants. At the end of the photoinhibitory period, F_v_/F_m_ was decreased in both WT and mutant plants. However, the decrease was significantly greater in the mutant lines, suggesting that the mutant plants are more sensitive to photoinhibition than the WT plants. After two hours of recovery period the F_v_/F_m_ value was similar in the WT and mutant plants and returned to the value of the initial values, suggesting full regeneration of PSII (Fig. 3).

**Figure 2.**
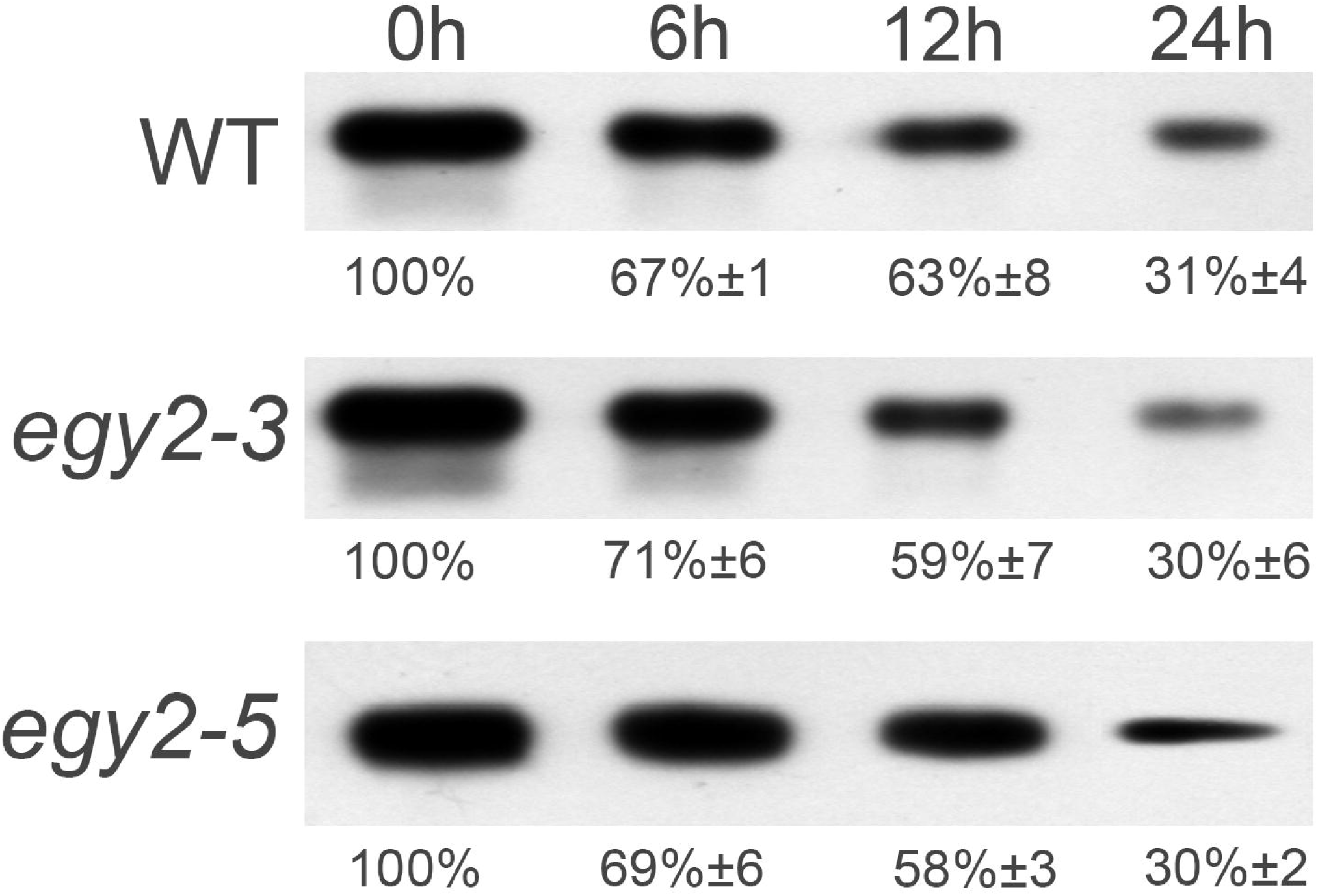
The comparison of rosette area of wild-type (WT), *egy2-3* and *egy2-5* lines. The rosette area was determined from plant photographs using ImageJ 1.50i software (National Institutes of Health, Bethesda, MD). The values shown the means ± SD determined by analysis of 90 plants representing three independent biological replicates.

**Figure 3.**
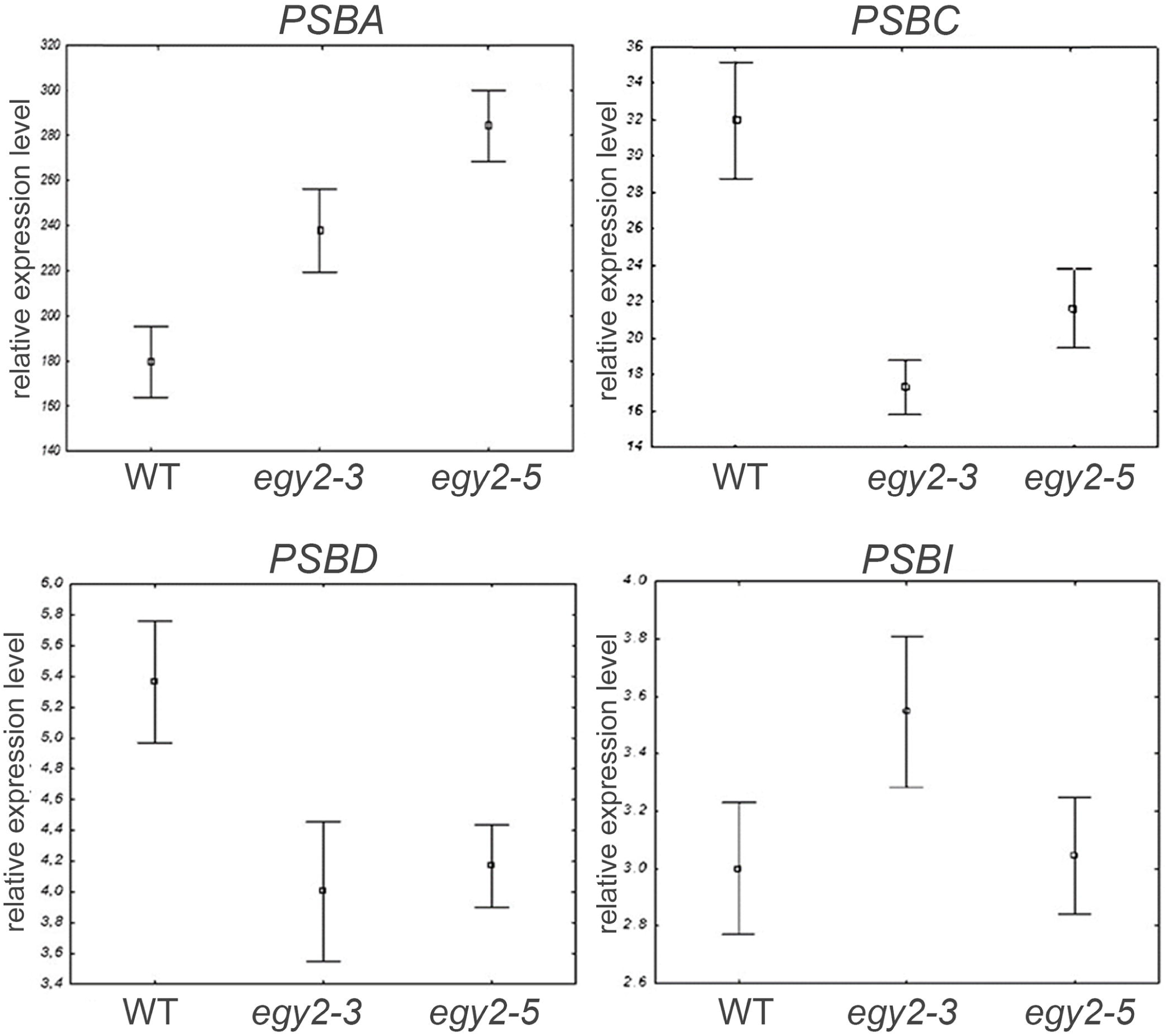
Maximum quantum efficiency of photosystem II photochemistry (Fv/Fm) in wild-type (WT), *egy2-3* and *egy2-5* leaves. The WT and mutant plants (black bars) were exposed for 2 h to 800 µmol m^-2^ s^-1^ (dark grey bars) followed by 2 h recovery at normal irradiance (pale grey bars). The values shown are the means ± SD determined by analysis of the Fv/Fm values of leaves representing three biological replicates (30 plants each).

### Abundance of selected PSII apoproteins in *egy2-3* and *egy2-5* mutants

Immunoblot analysis of the selected PSII apoproteins revealed no significant changes in the accumulation levels of the nuclear-encoded apoproteins Lhcb1-Lhcb6 (Fig. 4). However, three of the four analysed chloroplast-encoded apoproteins display altered accumulation levels in both mutant lines relative to their levels in WT plants. Altered accumulation levels were displayed by both apoproteins that form the PSII core centre, PsbA (D1) and PsbD (D2). However, the abundance of the PsbA apoprotein, which forms the reaction centre of PSII, was increased to approximately 150 % in both mutant lines, whereas a reduction in the accumulation level of the PsbD apoprotein to 67 % of the normal value was observed. A very similar decrease in abundance was observed for PsbC, which is an apoprotein associated with the CP43 complex, an inner PSII antennae. PsbC level decreased to 68 % in *egy2-3* and to 67 % in *egy2-5*. The amount of PsbI apoprotein, which is crucial for the stability of PSII supercomplexes, was unchanged in the mutant lines (Fig. 4).

**Figure 4.**
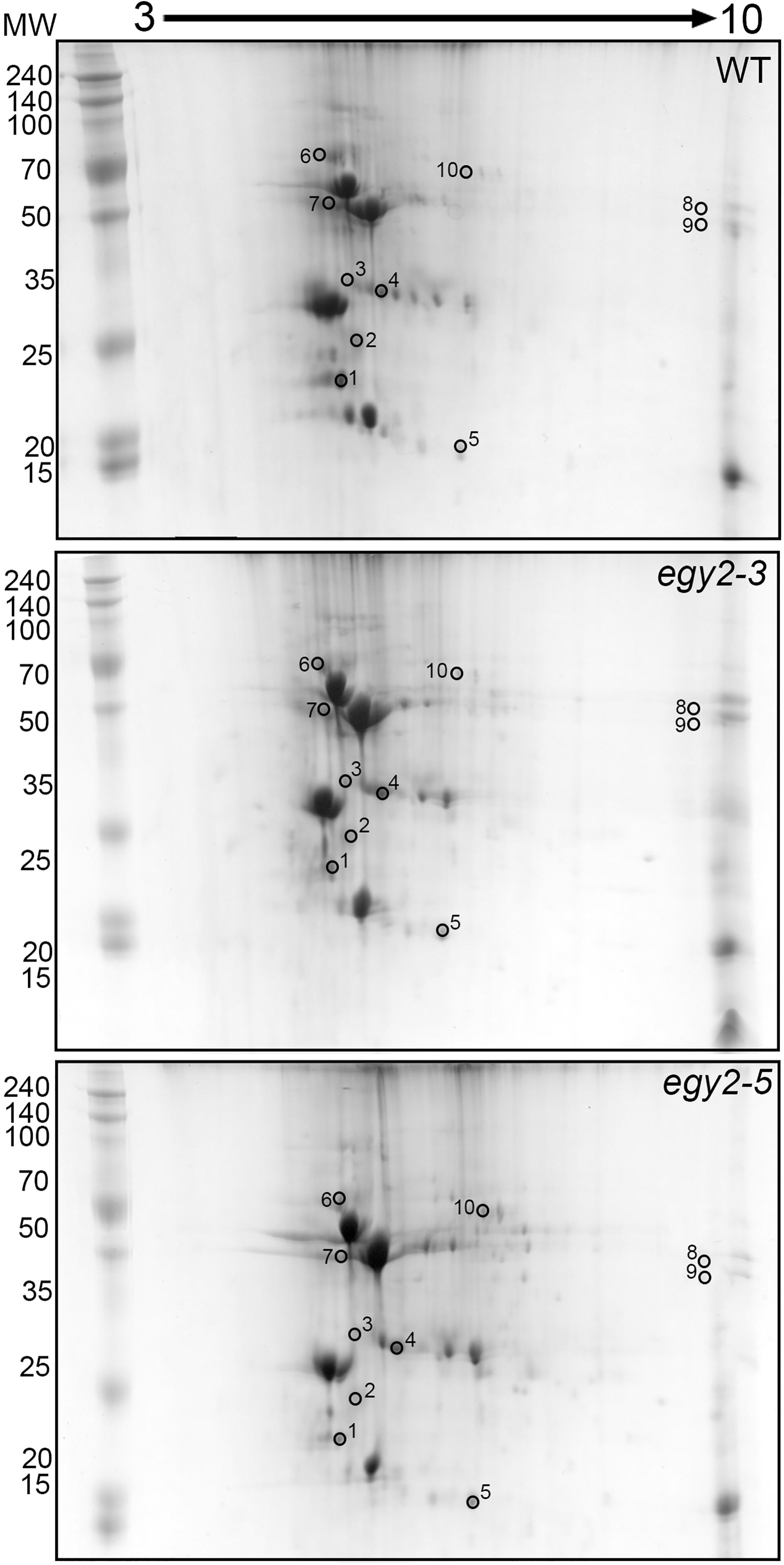
Immunoblot analysis of the levels of Lhcb1-6, PsbA, PsbC, PsbD and PsbI in wild-type (WT), *egy2-3* and *egy2-5* mutants. Total protein (2 µg) from each sample were subjected to immunoblotting analysis with specific primary antibodies. Quantification of the blots was performed using GelixOne software. The individual apoprotein content of the mutants was quantified as a percentage of the antibody signal strength in the WT (100%). “±” indicates the SD calculated from the analysis of samples from the four biological replicates. The asterisks indicate statistically significant differences between the WT and individual mutants.

To determine whether the increased PsbA accumulation level may be partially a result of impaired degradation, changes in PsbA abundance under high light conditions in mutant lines and WT plants were investigated. Due to the higher initial level of PsbA in the *egy2* mutant lines, the protein was present at a significantly higher level (approximately 150 % of its abundance in WT plants) throughout the whole period of plants exposure to 800 µmol m^-2^ s^-1^, however the percentage decrease in the protein abundance was similar in all analysed lines, both *egy2* mutants and WT (Fig. 5).

**Figure 5.**
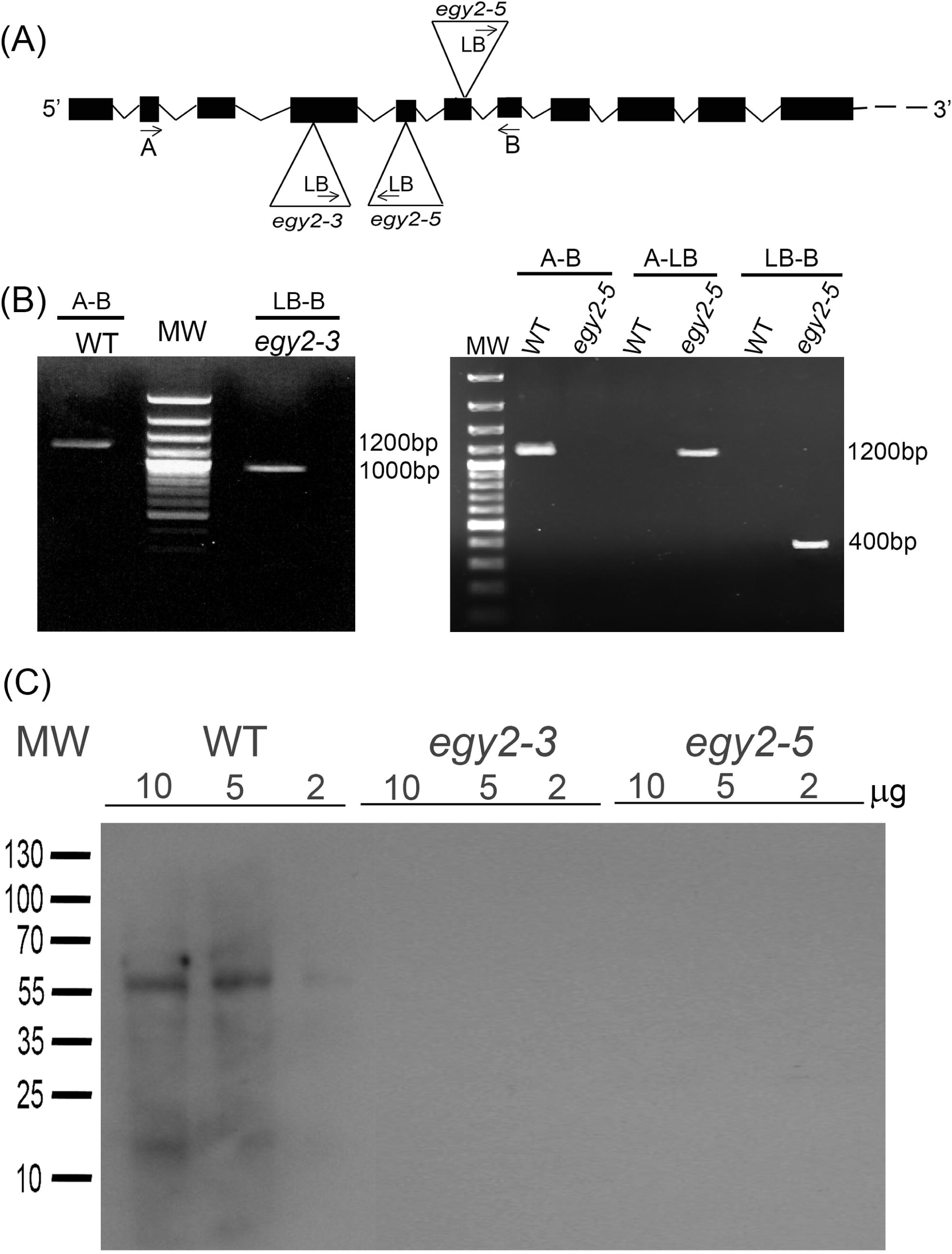
Immunoblot quantification of PsbA apoprotein in wild-type (WT), *egy2-3* and *egy2-5* mutant plants under high light conditions. Plants were exposed to 800 µmol m^-2^ s^-1^ for 6, 12 and 24 hours. Total protein (2µg) were immunologically analysed using an anti-PsbA antibody. GelixOne software was used to quantify the PsbA content. “±” indicates the SD determined in the analysis of samples obtained from three biological replicates, each of which was obtained by isolation of total protein from a minimum of 30 plants.

### qPCR analysis

To test the hypothesis that the observed changes in the abundance of chloroplast-encoded proteins are a consequence of changes in gene expression levels, real-time PCR was performed. The results revealed that the observed aberrations in PSII protein abundance in *egy2* mutants correlate strongly with changes in the transcription levels of the genes encoding these proteins. Significant increases in *PSBA* transcription to 132 % and 150 % of the levels seen in WT plants were observed in the *egy2-3* and *egy2-5* mutant lines, respectively. The abundance of *PSBD* transcripts was reduced to 74 % of that in the WT in the *egy2-3* mutant line and to 77 % of that in the WT in the *egy2-5 line*. A slightly greater reduction was observed in the accumulation of the PSBC transcript. In the *egy2-3* mutant line, the PSBC transcript level abundance was reduced to 54 % of that found in WT plants. In the *egy2-5* mutant line, the accumulation of *PSBC* transcripts corresponded to 67 % of the level found in WT plants. No statistically significant changes in the abundance of the *PSBI* transcript were observed (Fig. 6).

**Figure 6.**
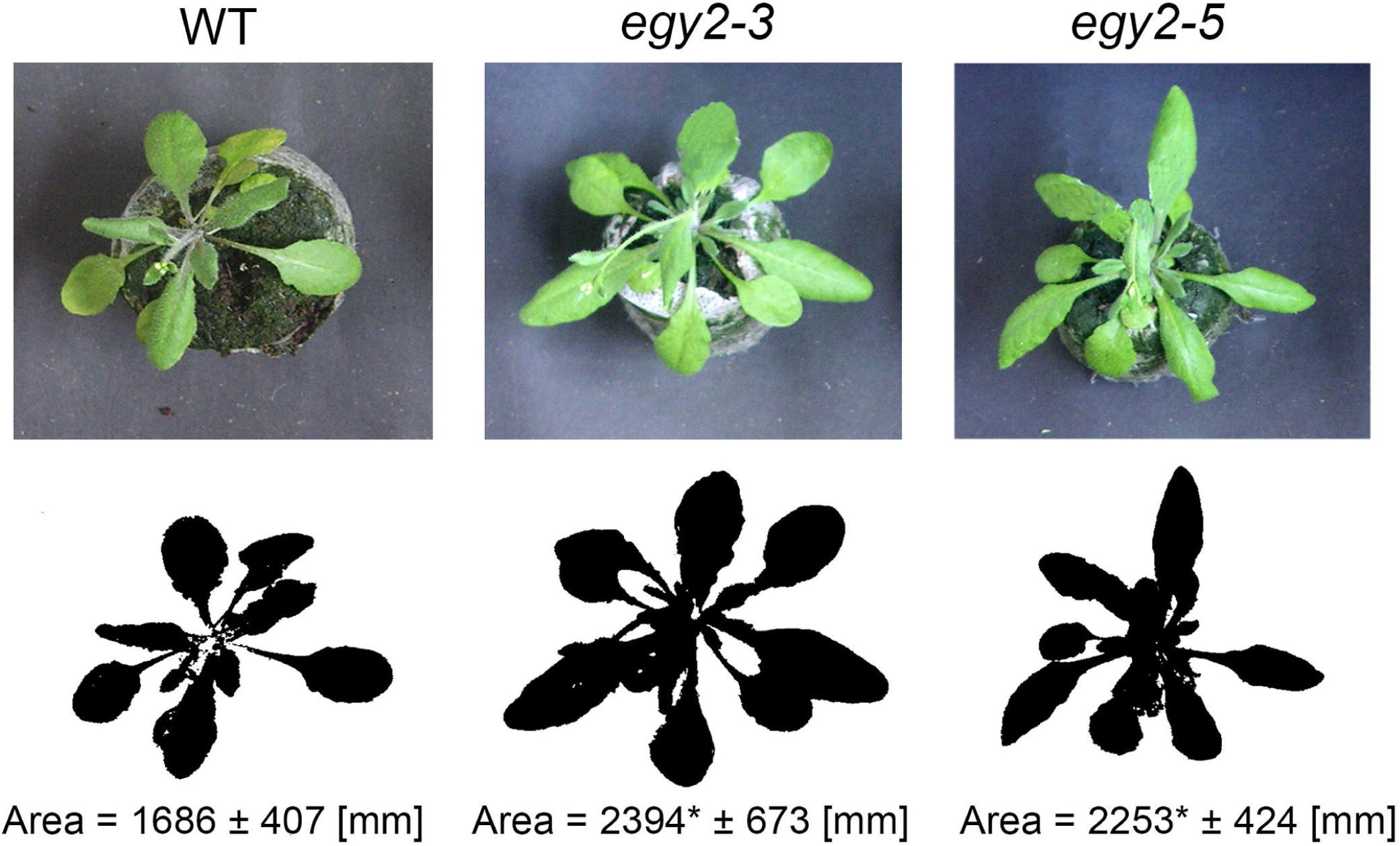
Relative expression levels of *PSBA, PSBC, PSBD* and *PSBI* in wild-type (WT) and *egy2* mutant (*egy2-3* and *egy2-5*) plants. Five micrograms of total RNA isolated from wild-type and *egy2* mutant plants were used for reverse transcription with random hexamers as primers, and 1 µl of cDNA was used in the qPCR reaction. The chloroplast ribosomal protein L2 gene was used as a reference. The results shown represent the means and standard errors determined by the analysis of samples from six biological replicates.

### Identification of proteins potentially involved in EGY2 dependent RIP

The EGY2 is located in thylakoid membrane and belongs to a group of intramembrane proteases, which are considered to activate membrane anchored transcription factors through proteolytic cleavage which release them form the membrane. To indicate potential transcription factors that may participate in EGY2 – dependent regulation of expression we performed comparative proteome analysis of thylakoid membranes. Two-dimensional electrophoresis was applied and protein spots whose abundance was increased at least 2-fold in both mutant lines were identified (Fig. 7). Only protein spots, the abundance of which was increased in two separate experiments, were selected for further analysis by LC - MS/MS. Selected spots were cut from two separate gels representing different *egy2* lines. For each sample LC - MS/MS analysis was performed and only proteins identified in both replicates were taken for further consideration. In 10 selected spots 218 proteins were identified by LC - MS/MS method (Fig. 7). The proteins with high score were screened for presence of plastid transcriptionally active chromosome proteins (pTAC) or other proteins related to chloroplast transcriptional machinery. pTAC10, FLN1 and pTAC16 were identified in 6th, 7th and 8th protein spot, respectively (Tab. 2). These proteins are known to be associated with PEP-mediated transcription of chloroplast genes (Arsova et al., 2010; Ingelsson and Vener, 2012; Chang et al., 2017).

**Figure 7.**
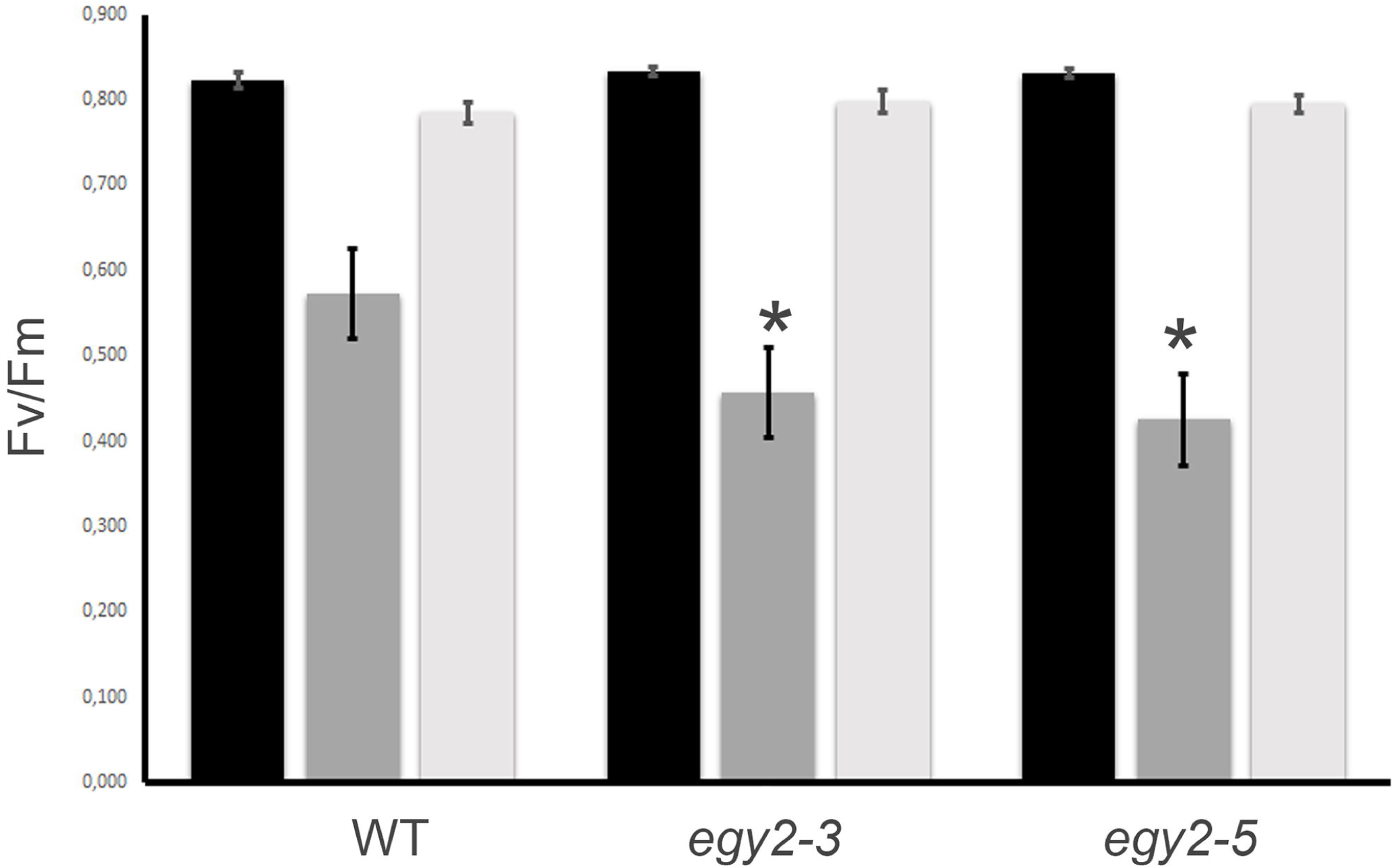
Two-dimensional electrophoresis gels. Thylakoid membrane proteins (150µg) isolated from wild type (WT), *egy2-3* and *egy2-5* lines were separated using 2-D gel electrophoresis with IEF (pH 3-10) and detected with Coomassie Brilliant Blue staining. Samples of proteins were obtained from four biological replicates of each plant lines. Ten protein spots were chosen for identification. All selected spots showed at least 2-fold up-regulation in comparison to WT, in both *egy2-3* and *egy2-5* mutant lines.

## Discussion

In the *egy2-3* mutant line, T-DNA insertion in the fourth exon and the homozygosity of its insertion were confirmed. In the *egy2-5* mutant line, the presence of two T-DNA insertions, one in the fifth and one in the sixth exon, was revealed; the homozygosity of that mutant line was also confirmed. In both mutant lines, the EGY2 protein was undetectable.

The analysis of rosette growth indicated that the final area of the rosette leaves is significantly larger in both *egy2* mutant lines than in WT plants. In both *egy2* mutant lines, slight but statistically significant decreases in chlorophyll and carotenoid content were also observed, as well as a reduction in the chlorophyll *a/b* ratio. This result is inconsistent with previous reports that *egy2* mutants do not exhibit changes in chlorophyll content (Chen et al., 2012). The diverse results may be associated with plant growth conditions and with the stage of plant development at which the measurements were performed. The chlorophyll measurements performed by Chen were conducted on “adult plants” grown under constant white light 100 µmol m^-2^ s^-1^ (Lu et al., 2002; Chen et al., 2012) while our analysis were carried out on plants with first flower open grown under long-day conditions (16 h of light and 8 h of darkness) at slightly higher irradiance of 110 µmol m^-2^ s^-1^.

The lack of EGY2 protease leads to a significant increase in PsbA accumulation and to significant reductions in the abundance of PsbC and PsbD proteins. Altered protein levels are in turn reflected in some parameters that are related to PSII functioning in both normal and high irradiance conditions. The PAM fluorescence measurements of chl a measured on plants growing in normal light conditions revealed increased non – photochemical quenching (NPQ) and minimum fluorescence yield (F_0_) in both of the *egy2* mutant lines. These two parameters are used to quantify non – photochemical quenching (Maxwell and Johnson, 2000). NPQ is a process in which excess of absorbed light energy is dissipated into heat. The process can be triggered directly by protonation of antenna components or indirectly by the xanthophyll cycle and involves three key elements: the LHCII antenna, violaxanthin de-epoxidase and the PsbS protein. The minor peripheral antenna CP26 (Lhcb5) and CP29 (Lhcb4) were also proven to be enriched in xanthophyll cycle carotenoids and displayed high level of quenching (Bassi and Caffarri, 2000). The monomer peripheral antennae are as well consider as the site for NPQ (Ahn et al., 2008; Avenson et al., 2009). The protein component of CP43 (PsbC), acts as an internal PSII energy antenna, transmitting the excitation energy of electrons from the external antennae to the PSII reaction centre. In addition, together with Lhcb5, PsbC plays an important role in maintaining the strong affinity between peripheral antenna and the PSII core and in docking LHCII to the PSII core at so-called S-sites of PSII-LHCII supercomplexes (Boekema et al., 1999; Caffarri et al., 2009). Decreased PsbC content in *egy2* mutant lines can lead to increased pool of free LHCII timers and the decrease in excitation energy transfer from the antennae via PsbC to the PSII reaction centre. The excess excitation energy within the peripheral antennas has to be dissipated thermally, and this is manifested in an increase in the NPQ parameter. The F_0_ parameter, in turn, is altered by D1 damage but not by the xanthophyll cycle (Murchie and Lawson, 2013). It is possible to interpret the increase in the F_0_ parameter as a reduction of the rate constant of energy trapping by PSII centres (Havaux, 1993), which could also result from a physical dissociation of LHCII from the PSII core. Such an effect was previously observed in several plant species following heat damage (Armond et al., 1980). In this light, the observed increase in F_0_ may also be due to the reduced PsbC content, which causes elevated dissociation of LHCII from the PSII core. To investigate how the changes in chlorophyll fluorescence and variations in PsbA/D and PsbC stoichiometry influence PSII activity under photoinhibitory conditions, changes in PsbA level and the sensitivity of *egy2* mutants to photoinhibition were measured. The somewhat more marked decrease in F_v_/F_m_ under photoinhibitory conditions was observed, which indicates a slightly higher sensitivity of *egy2* mutants to photoinhibition, is likely to result from the disturbed stoichiometry of PsbA/PsbD. However, the recovery of PSII efficiency after the termination of photoinhibitory conditions was shown to be faster in the *egy2* mutants than in WT plants. This phenomenon may be partly because of the higher initial PsbA level. The mutant plants consistently maintain a significantly larger PsbA pool than the WT plants. The observed changes in the PsbA content during exposure of plants to high irradiance indicate, however, that the lack of EGY2 does not visibly impairs the rate of degradation of PsbA and the increased PsbA abundance is rather a consequence of increased synthesis than impaired degradation. From a physiological point of view, an interesting fact is that the genes encoding the PSII reaction center proteins are counter-regulated. This situation was also observed in other experimental conditions like 7 days cold treatment (4°C), treatment with DMTU, which is H_2_O_2_ scavenger or overexpression of ABA responsive ABF3 transcription factor (Hruz et al., 2008). The increase in *PSBA* gene expression with simultaneous decrease in *PSBD* and *PSBC* was observed also in double mutant in calmoduling binding transcription activator *camta1 camta 2*, drought treatment of double mutant in SNF1-related protein kinases 2 *srk2cf* or triple mutant in cytokinin receptor *ahk2/ahk3/ahk4* and several other experimental conditions. (Hruz et al., 2008). Within the chloroplast genome, *PSBD* and *PSBC* genes are located in a single operon, whereas the *PSBA* gene is located on a separate one, however, both of them are classified as class I operons, transcribed by plastid-encoded plastid RNA polymerase (PEP; Hajdukiewicz et al., 1997). The promoter specificity of PEP is achieved by the sigma factors. In *Arabidopsis thaliana* six sigma factors (SIG1-6) have been identified. The *PSBC/PSBD* promoter was found to be recognized only by SIG5 sigma factor while SIG1, SIG2 and SIG5 factors were able to bind to *PSBA* operon (Chi et al., 2015). The counter-regulation of *PSBA* and *PSBC/PSBD* operons can therefore be a consequence of recognition by different types of PEP-complexes, transcription fine tuning by diverse PEP-associated proteins (PAPs) or the resultant of both of these phenomena. In two – dimensional electrophoresis and LC - MS/MS analysis three PAPs accumulating in thylakoid membranes of *egy2* mutants were identified, namely FLN1, pTAC10 and pTAC16. The role of these proteins in regulation of chloroplast genes transcription is, however, poorly investigated. Nonetheless the FLN1 was found to interact with another PAPs – thioredoxin z (TRX z) in a thiol-dependent way, indicating on redox–dependent transcriptional gene regulation pathway (Arsova et al., 2010, Wimmelbacher and Börnke, 2014). pTAC10 was also found to interact with TRX z, however the mechanism of this interaction remains elusive (Chang et al., 2017). Moreover, pTAC10 was proven to play crucial role in the proper assembly of the PEP complex and chloroplast development (Chang et al., 2017). The role of pTAC16 in regulation of chloroplast gene expression remains unknown. The protein was found to associate with nucleoid regions but is not essential for its formation and composition (Ingelsson and Vener, 2012). The above results suggest that FLN1, pTAC10 and pTAC16 may participate in EGY2 – dependent regulated intramembrane proteolysis process.

## Author Contribution

MA: Developing of the article concept, performing of experiments and data analysis, drafting the article,

LM: selection and basic analysis of the homozygous mutant lines

EK: participation in the Real-Time experiments

EPL: developing the method of isolation of the thylakoid membrane proteins for IEF separation.

RL: participation in the design and realization of experiments. Participation in the development of the concept of the work, participation in 2D experiments. Head of the group.

## Founding

This work was supported by the Polish National Science Center based on decision number DEC-2014/15/B/NZ3/00412.

## Acknowledgements

The equipment used for LC - MS/MS analysis was sponsored in part by the Centre for Preclinical Research and Technology (CePT), a project co-sponsored by European Regional Development Fund and Innovative Economy, The National Cohesion Strategy of Poland. Ewelina Paluch-Lubawa is Adam Mickiewicz University Foundation scholar in 2017/2018 academic year

---

**Table 1.** Comparison of the rosette size, chloroplast pigment content of leaves and chlorophyll fluorescence parameters in wild-type (WT) plants and *egy2* (*egy2-3* and *egy2-5*) mutants in normal light conditions. “±” indicates the SD calculated from the analysis of three biological replicates (20 plants each). * - indicate statistically significant differences between the WT and individual mutants, ^a^ - indicate statistically significant differences between *egy2-3* and *egy2-5.*

|  | WT | *egy2-3* | *egy2-5* |
| --- | --- | --- | --- |
| rosette size [mm] | 1686 ± 407 | 2394* ± 673 | 2253*± 424 |
| chl a | 528 ± 86 | 418* ± 54 | 451*± 71 |
| chl b | 201 ± 37 | 164* ± 20 | 176*± 26 |
| chl a/chl b ratio | 2.6± 0.01 | 2.5*± 0.01 | 2.5* ± 0.01 |
| carotenoids | 114 ± 18 | 95* ± 11 | 101*± 14 |
| F<sub>0</sub> | 138 ± 8 | 190*± 19 | 250*<sup>a</sup> ± 25 |
| F<sub>v</sub>/F<sub>m</sub> | 0.844± 0.01 | 0.838 ± 0.01 | 0.842 ± 0.00 |
| F<sub>v</sub>'/F<sub>m</sub>' | 0.755± 0.02 | 0.762 ± 0.02 | 0.754 ± 0.02 |
| NPQ | 0.212± 0.08 | 0.328*<sup>a</sup> ± 0.06 | 0.371*<sup>a</sup> ± 0.06 |
| qP | 0.892± 0.01 | 0.894 ± 0.03 | 0.883 ± 0.03 |
| ΦPSII | 0.673± 0.02 | 0.681 ± 0.04 | 0.665 ± 0.03 |

**Table 2.** Elements of chloroplast transcriptional machinery identified in protein spots with increased accumulation level in *egy2* mutants. I, II-indicates separate LC-MS/MS analysis.

| Locus | protein name | spot number | MW (molecular weight) | pI (isoelectric point) | protein score |  | number of peptid mashes |  | protein coverage (%) |  |
| --- | --- | --- | --- | --- | --- | --- | --- | --- | --- | --- |
|  |  |  | theor/exper | theor/exper | I | II | I | II | I | II |
| At3g48500 | PTAC10 | 6 | 79/ 79,2 | 5,0/4,9 | 185 | 352 | 6 | 12 | 7,3 | 17 |
| AT3G46780 | PTAC16 | 8 | 54/54,3 | 8,9/8,4 | 720 | 537 | 14 | 10 | 26,5 | 20 |
| AT3G54090 | FLN 1 | 7 | 54/54,8 | 5,8/5,4 | 166 | 334 | 4 | 7 | 9,6 | 18 |

