## Supplementary figures and images for "*Arabidopsis thaliana egy2* mutants display altered expression level of genes encoding crucial photosystem II proteins"

### Supplementary Materials

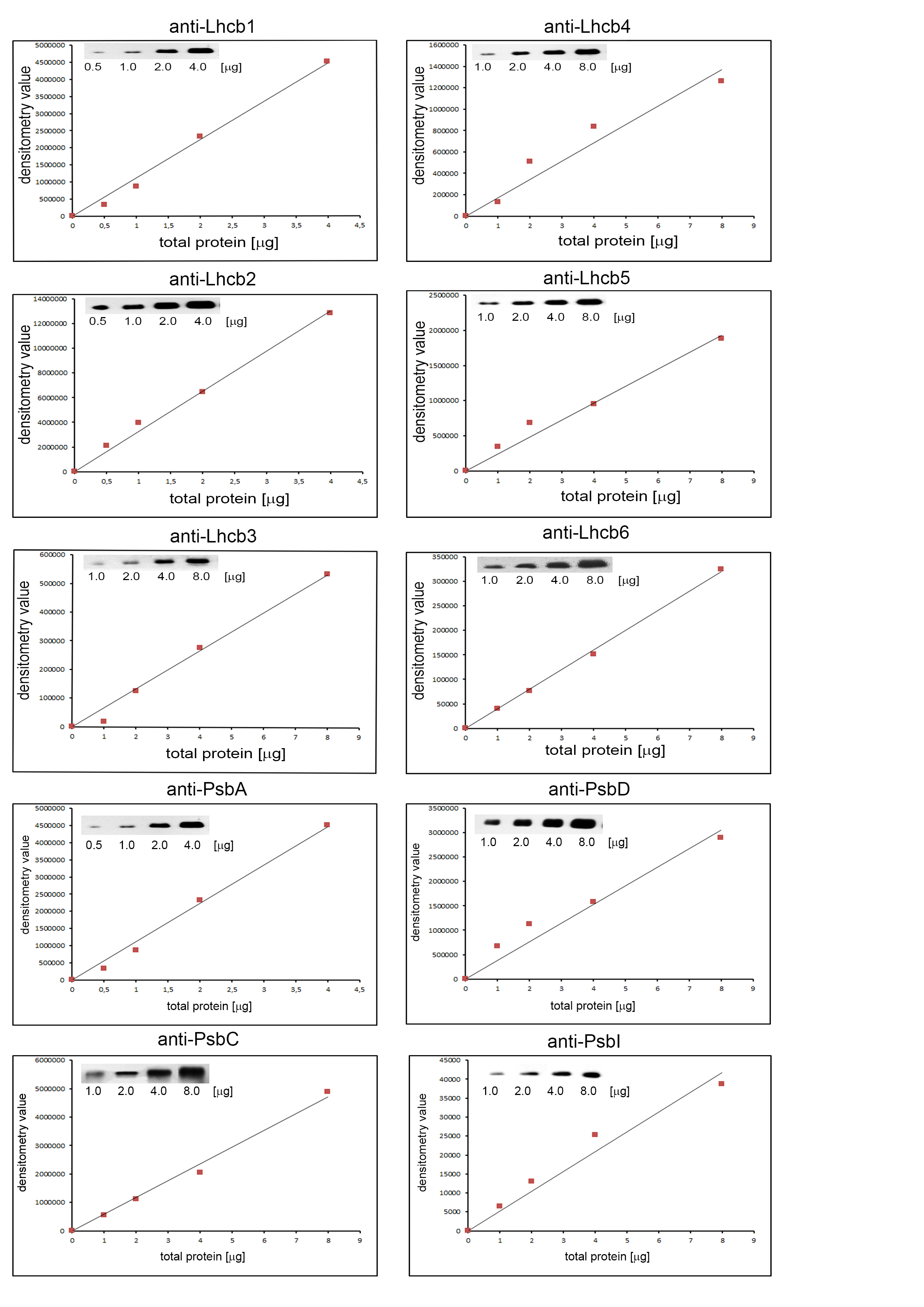
